# Conformity to Bergmann’s rule in birds depends on nest design and migration

**DOI:** 10.1101/686972

**Authors:** Mark C. Mainwaring, Sally E. Street

## Abstract

Species’ geographic ranges and range limits are thought to be determined by climate, and across climatic gradients the morphology of populations varies non-randomly. Ecogeographic rules seek to characterise such variation, with Bergmann’s rule positing that organisms inhabiting colder environments are typically larger-bodied than those inhabiting warmer environments. While Bergmann’s rule has been supported across a range of taxonomic groups, how organisms’ behaviour may moderate its effect remains unclear. Here we investigate whether conformity to Bergmann’s rule among birds of the Western Palearctic varies in relation to nest design and migratory behaviour, using phylogenetic comparative analyses. We test predictions using data on nest structure and location, migration, body mass, latitudinal distribution, annual mean temperature and phylogenetic relatedness for a sample of >500 species. We find that conformity to Bergmann’s rule depends strongly on migratory behaviour: non-migratory species breeding at colder, more northerly latitudes are larger-bodied, while body mass is unaffected by climate in short- and long-distance migrants. Among non-migratory species, conformity to Bergmann’s rule depends, further, on nest design: species with more open nests, who are thus most exposed to adverse climatic conditions while breeding, conform most strongly to Bergmann’s rule. Our findings suggest that enclosed nesting and migration allow smaller bodied species to breed in colder environments than their body size would otherwise allow. Therefore, we conclude that organisms’ behaviour can strongly affect exposure to environmental selection pressures.

## Introduction

Species’ geographic ranges and range limits are influenced by abiotic factors such as environmental conditions and physical barriers to dispersal, and by biotic factors such as competition for resources and the risk of predation (Gaston 2003, 2009). Whilst the relative importance of these factors varies between species and geographic regions, the availability of ambient and productive energy primarily determines species’ geographic ranges and range limits (Currie et al. 2004). This is because species’ range edges occur at climatic thresholds and vary predictably with changes in climatic conditions (Gaston 2003, 2009), because organisms have a unique set of environmental conditions in which they can survive and reproduce (Blackburn and Gaston 1996). The morphology of individuals also varies non-randomly across geographic clines. Ecogeographic principles seek to characterise such variation, with Bergmann’s rule being one of the most widely known and enduring examples (James, 1970). Bergmann’s rule posits that species inhabiting colder environments are larger than species inhabiting warmer environments, because larger bodied organisms have a lower surface area to volume ratio and thus a greater ability to conserve heat (Bergmann 1847).

However, evidence that body size and morphology correlate in accordance with Bergmann’s rule remains variable, and its general validity across diverse animal taxa has been questioned (Blackburn et al. 1999, Meiri and Dayan 2003). At the intraspecific level, a substantial minority (∼25-30%) of bird species fail to conform to Bergmann’s rule (Ashton 2002, Meiri and Dayan 2003). While Bergmann’s rule is broadly supported across bird species globally, the magnitude and direction of relationships between body size and climate varies widely among geographic regions and taxonomic groups (Olson et al. 2009). Further, within taxonomic subsets, Bergmann’s rule is not supported in the majority of bird genera, families and orders (Olson et al. 2009). Taken together, these results suggest that while broadly valid, conformity to Bergmann’s rule is mediated by other factors.

Organisms are able to buffer against environmental selection pressures to some extent via their behaviour (Odling-Smee et al. 2003). Nesting behaviour may be particularly important in this regard because nests are structures built to contain and protect offspring and the attending parents during the breeding season (Martin et al. 2017, Hansell 2000). Both aspects of nest structure and location can affect the exposure of parents and offspring to climatic conditions (Biancucci and Martin 2010). However, the potential relationship of nests to species’ responses to environmental conditions such as those predicted by Bergmann’s rule is currently underexplored. In particular, incubating parents and broods of species with open nests may be more exposed to the adverse effects of cold weather than those of species with enclosed nests (Mainwaring et al. 2017). We therefore predict that species with enclosed nests should not conform as strongly to Bergmann’s rule as open nesting species. This prediction has not, so far, found support at the intraspecific level: body mass is no more likely to increase in colder climates within species laying in open nests than within those with enclosed nests (including tree cavities, burrows and domed nests built of vegetation, Meiri and Dayan 2003). Bergmann’s rule has been proposed, rather, to depend on migratory behaviour, as migrating species avoid extreme winter temperatures that may select for large body size at high latitudes (Ashton 2002, Meiri and Dayan 2003). In support of this idea, some analyses find that body mass is more likely to increase in colder temperatures within sedentary rather than migratory bird species, but this pattern is not consistent across studies using different statistical approaches (Meiri and Dayan 2003, Ashton 2002). As yet, the potential role of nesting and migration in mediating Bergmann’s rule across species, where variation in body mass and geographic range far exceeds that within species, has not yet been investigated.

To investigate the potential role of nest design and migration as mediators of conformity to Bergmann’s rule, we examined the combined effects of nest type, migration and climatic conditions on body mass among the bird species of the Western Palearctic. We focus on this region due to dramatic variation in environmental conditions over large-scale latitudinal gradients in the Northern hemisphere, in which ambient temperatures are considerably colder in northerly than southerly regions. We test predictions with phylogenetic comparative analyses based on a sample of >500 species with data on nest design, migratory behaviour, breeding latitude and breeding range temperature. We fit models in which the slope of body mass on latitude or temperature is allowed to vary between species with differing nest designs and migratory strategies. We predict that conformity to Bergmann’s rule should depend on nest design as follows: body mass should increase with latitude and decrease with temperature most strongly among species with open nests, followed by those with semi-open nests while those with enclosed nests conform least to Bergmann’s rule. In terms of migration, we predict the strongest conformity to Bergmann’s rule among sedentary (non-migrating) species, followed by short-distance and long-distance migrants. If nest type and migration have equally important effects on conformity to Bergmann’s rule, we should find the same effects of nest design within each category of migratory behaviour, and vice versa. Alternatively, if either nest design or migration is a more important mediator of conformity to Bergmann’s rule, one should override the effect of the other.

## Materials and methods

### Study species

We examined relationships between nest design, migration, body mass and breeding climate among bird species listed as breeding residents in the Birds of the Western Palearctic book series (Cramp and Simmons 1977, 1980, 1983, Cramp 1985, 1988, 1992, Cramp and Perrins 1993, 1994a, 1994b). Data on nest structure and location (categorised based on descriptive information), body mass, latitudinal range and migratory behaviour were collected from the same source for n=857 species, of which n=769 could be matched with the Jetz et al. (2012) molecular phylogeny. Bioclimatic data (annual mean temperature, ‘BIO1’) were obtained from the WorldClim global climate database (Fick and Hijmans 2017) and matched to species’ ranges using distribution maps from BirdLife International (BirdLife International & Handbook of the Birds of the World 2018). Of species included in the phylogeny, data on nest structure and location were available for n=538 species, body mass for n=518 species, latitudinal range for n=530 species and migratory behaviour for n=538 species. We obtained bioclimatic data only for species that were included in the phylogeny and which had available body mass, nest design and migration data (n=518). 3 such species lack temperature data either because the species is no longer recognized in the latest HBW-BirdLife Taxonomic Checklist (version 3.0, November 2018) or because the species’ range is too small to be matched to bioclimatic data at the chosen grid resolution, leaving n=515 with annual mean temperature data. The full dataset used for analysis, along with associated R code and additional relevant files, is available in the Supporting Information.

### Classifying nest types

Here, we consider nest type to be more complex than simply the structure of the nest itself, because the location in which the nest is built also strongly affects its exposure to climatic conditions (Mainwaring et al. 2014). For example, open cup-shaped nests built in vegetation should be more exposed to environmental conditions than open cup-shaped nests built inside tree cavities (von Haartman 1957). Therefore, we combine both aspects of the structure and location of birds’ nests to produce an appropriate single nest design factor as follows. Nest structure is classified following Hansell (2000) in figure 3.2 as either cup, plate, scrape, bed, dome, dome and tube or burrow, while nest location is classified as open, semi-open or enclosed in which open refers to fully exposed nest sites (such as waders nesting on bare ground), semi-open refers to those nests that are largely concealed from all sides by, for example, being located in dense vegetation and enclosed refers to nests in tree cavities and alike (von Haartman 1957, Alerstam and Hogstedt 1981, Hansell 2000). We then combine information on nest structure and location to classify species’ overall nest type as either open, semi-open or enclosed (Table S1). We consider as ‘open’ nest types only open nest structures (cup, plate, scrape or bed nests) built in open locations. We consider as ‘enclosed’ nests both nests of any structure located inside cavities, and enclosed nest structures (dome, dome and tube or burrow) built in any location. Finally, we treat open nest structures (cup, plate, scrape or bed nests) built in ‘semi-open’ locations as ‘semi-open’ nests, an intermediate state between fully open and fully enclosed nest designs. In Supporting Information, we present the results of additional analyses treating nest structure and location separately.

### Quantifying breeding climate

Bergmann’s rule is generally tested using measures of climatic conditions across a species’ entire range. However, since our predictions concern how body mass may be affected by exposure to climatic conditions while breeding, here we investigate relationships between species’ body mass and climatic conditions of the breeding range specifically. We used two variables to capture climatic conditions in the breeding range: breeding range latitude and breeding range temperature. We obtained northernmost and southernmost latitudes of the breeding ranges for each species from distribution maps in the Birds of the Western Palearctic book series (Cramp and Simmons 1977, 1980, 1983, Cramp 1985, 1988, 1992, Cramp and Perrins 1993, 1994a, 1994b). For analyses, we used a single latitudinal measure, ‘breeding latitude midpoint’, taken as the mean of the northernmost and southernmost breeding latitudes. To estimate breeding range temperature, we matched annual mean temperature data from WorldClim (Fick and Hijmans 2017) to species’ ranges from BirdLife International (BirdLife International & Handbook of the Birds of the World 2018) using functions from the R packages ‘rgdal’ (Bivand et al. 2018) and ‘letsR’ (Vilela and Villalobos 2015). Species’ ranges were converted to presence-absence matrices with 0.5 degree grid cell resolution (∼55km at the equator), counting a species as present if its range covered 10% or more of a cell. We exclude uncertain records (presence codes 2 = ‘probably extant’, 3 = ‘possibly extant’ and 6 = ‘presence uncertain’) and records from outside of species’ native ranges (all except origin code 1 = ‘native’ and 2 = ‘reintroduced’). To limit records to ranges in which birds may breed, we select only records from the birds’ resident or breeding season ranges (season codes 1 = ‘resident’ or 2 = ‘breeding season’), thereby excluding non-breeding season and passage ranges, and records of uncertain seasonality (season codes 3 = ‘non-breeding season’, 4 = ‘passage’ and 5 ‘seasonal occurrence uncertain’). We obtained annual mean temperature data at 10 minutes of a degree resolution, matching it to each grid cell where a species is present. Since the climatic data is higher resolution than the presence absence matrix, we average climatic data across cells at the coarser 0.5-degree resolution to match the presence absence matrix. Finally, we summarise breeding range temperature at the species-level by taking the mean of annual mean temperature across all occupied cells in each species’ presence-absence matrix.

### Quantifying body sizes

We measured species’ body sizes as the mean body mass of males and females during the breeding season, preferring estimates from the UK (where appropriate) due to larger sample sizes, using data from the Birds of the Western Palearctic (Cramp and Simmons 1977, 1980, 1983, Cramp 1985, 1988, 1992, Cramp and Perrins 1993, 1994a, 1994b). Body masses of unknown sex were used where body mass was not reported separately for males and females, following e.g. Møller et al. (2010).

### Quantifying migratory behaviour

Using data from the Birds of the Western Palearctic (Cramp and Simmons 1977, 1980, 1983, Cramp 1985, 1988, 1992, Cramp and Perrins 1993, 1994a, 1994b), we categorised species’ migratory behaviour, distinguishing between sedentary (non-migratory) species, short-distance migrants and long-distance migrants. Sedentary species are those species that remain in the same area year-round and are thus residents, whilst short-distance migrants migrate south each autumn to over-winter either in southern Europe or northern Africa, and long-distance migrants migrate south each autumn to over-winter in sub-Saharan Africa.

### Statistical analyses

We test predictions using Bayesian phylogenetic generalized linear mixed models and phylogenetic generalized least squares regression, implemented in the MCMCglmm R package (Hadfield 2010) and BayesTraits software (Pagel 1999, Pagel et al. 2004, Meade and Pagel 2016) respectively. To test for Bergmann’s rule across the whole sample of species, we fit body mass as the outcome variable, predicted by either breeding range latitude or temperature. To investigate whether conformity to Bergmann’s rule is affected by nesting variables and migration, we include an interaction term allowing slopes of body mass on latitude or temperature to vary between species with different nest characteristics and/or migratory strategies. For models incorporating interactions, sample sizes are sufficient that there are at least 10 species for every slope estimated. Body mass is log10-transformed to correct for a strong positive skew, while breeding latitude and temperature are roughly normally distributed and are left untransformed.

Accounting for phylogenetic non-independence is essential in cross-species comparative analyses to avoid pseudoreplication and biased parameter estimates (Freckleton et al. 2002). We obtained trees from a comprehensive global bird phylogeny (Jetz et al. 2012), selecting a version constructed using only species with molecular data, based on the Hackett et al. (2008) ‘backbone’ phylogeny. For the majority of our analyses we use a single maximum clade credibility (MCC) phylogeny based on a posterior sample of 10,000 trees, created with TreeAnnotator (Drummond et al. 2012). However, to ensure analyses are robust to phylogenetic uncertainty, we repeat one of our main analyses across a posterior distribution of 3000 trees in BayesTraits (Pagel 1999, Meade and Pagel 2016). This approach uses MCMC to estimate model parameters across the posterior tree distribution, thereby incorporating both model and phylogenetic uncertainty into results (Pagel et al. 2004). We find qualitatively identical results, both when sampling trees in proportion to their likelihood and when visiting each tree for an equal number of iterations (Table S2). Therefore, we are confident that our results are not substantially affected by phylogenetic uncertainty. For MCMCglmm analyses we quantify the influence of phylogeny on results by estimating heritability (*h*^2^), the proportion of residual variance attributable to phylogenetic relationships equivalent to Pagel’s λ for PGLS regression (Hadfield and Nakagawa 2010). Like Pagel’s λ, *h*^2^ varies from 0, equivalent to an ordinary non-phylogenetic regression with a random error structure, to 1, where the covariation in residual errors is directly proportional to phylogenetic relationships, assuming a Brownian motion model of trait evolution (Pagel 1999, Freckleton et al. 2002).

For all models, we ran MCMC chains of sufficient length to obtain effective sample sizes of at least 1000 for all model parameters (MCMCglmm = 501,000 iterations, sampling every 100 iterations, with a burn-in period of 1000 iterations; BayesTraits = 5,050,000 iterations, sampling every 1000 iterations, with a burn-in of 50,000 iterations). For MCMCglmm analyses, we use default, diffuse normal priors for predictor variables (mean=0, variance=10^8^) and commonly used inverse-Wishart priors for the residual variance and phylogenetic random effect (with V=1, nu=0.002) (Hadfield 2019). For BayesTraits analyses, we use default minimally-informative, uniform prior distributions for all parameters, with a range of −100 to 100 for fixed effects and 0 to 1 for Pagel’s λ (Meade and Pagel 2016). For every model, we ensured that chains had converged on the posterior distribution, that burn-in periods were sufficient and that chains did not have problematic levels of autocorrelation by confirming sufficient effective sample sizes and by visual examination of chain plots. For all parameter estimates, we report means and 95% credible intervals from posterior distributions. Additionally, for each model we report R^2^ estimated as the squared Pearson’s correlation of the observed and fitted Y values.

### Data deposition

Data will be made available in the Dryad Digital Repository upon acceptance.

## Results

### Support for Bergmann’s rule

We find support for Bergmann’s rule across the whole sample: species’ body mass increased with breeding range latitude midpoint (*β* = 0.002 [<-0.001, 0.004], *n* = 513, *h*^*2*^ = 0.990 [0.973, 0.998], *R*^*2*^ = 0.026, Fig. S1a) and decreased with breeding range temperature (*β* = −0.003 [-0.005, <0.001], *n* = 515, *h*^*2*^= 0.989 [0.973, 0.998], *R*^*2*^ = 0.019, Fig. S1b). These relationships are, however, fairly weak, with breeding climate explaining less than 3% of the variation in body mass. Translating effects to the data scale, the model predicts that body mass increases from 277g to 363g across the range of breeding latitudes (16 to 79.5 degrees), while body mass decreases from 384g to 297g from the lowest to the highest breeding range temperatures (-16°C to 28°C).

### Effects of nest design and migration on conformity to Bergmann’s rule

When modelling interactions of nest design and breeding climate, we find that nest design affects conformity to Bergmann’s rule (Table 1). Specifically, in semi-open nesting species, we find similar effects to those found across the whole sample: body mass increases slightly with breeding range latitude and decreases with breeding range temperature. However, among open and enclosed nesting species, there is little to no relationship between body mass and either latitude or temperature. Repeating analyses for nest location and structure separately suggests that this pattern is driven primarily by nest location rather than structure (Tables S3, S4).

**Table 1.** Interaction of Bergmann’s rule with nest design.

a) Latitude
| Nest design | $\beta$ Estimates & 95% CI | $h^2$ & 95% CI | $n$ species | $R^2$ |
| --- | --- | --- | --- | --- |
| Open | <-0.001 [-0.003, 0.004] | 0.987 [0.968, 0.998] | 513 | 0.198 |
| Semi-open | 0.003 [-0.001, 0.006] |  |  |  |
| Enclosed | 0.002 [-0.003, 0.007] |  |  |  |

b) Temperature
| Nest design | $\beta$ Estimates & 95% CI | $h^2$ & 95% CI | $n$ species | $R^2$ |
| --- | --- | --- | --- | --- |
| Open | 0.001 [-0.003, 0.006] | 0.988 [0.970, 0.998] | 515 | 0.184 |
| Semi-open | -0.005 [-0.009, -0.001] |  |  |  |
| Enclosed | -0.004, -0.010, 0.002 |  |  |  |

$\beta$ estimates = mean regression slopes from posterior distributions for body mass on a) latitude and b) temperature, fitted for species with different nest designs, $h^2$ = mean heritability (phylogenetic signal) 95% CI = 95% credible intervals.

When including interaction terms for migration and breeding climate, we find support for the predicted effect of migration on conformity to Bergmann’s rule: among sedentary species, body mass increases with breeding latitude and decreases with breeding range temperature, but not in short- or long-distance migrants (Table 2, Fig. 1).

**Table 2.**
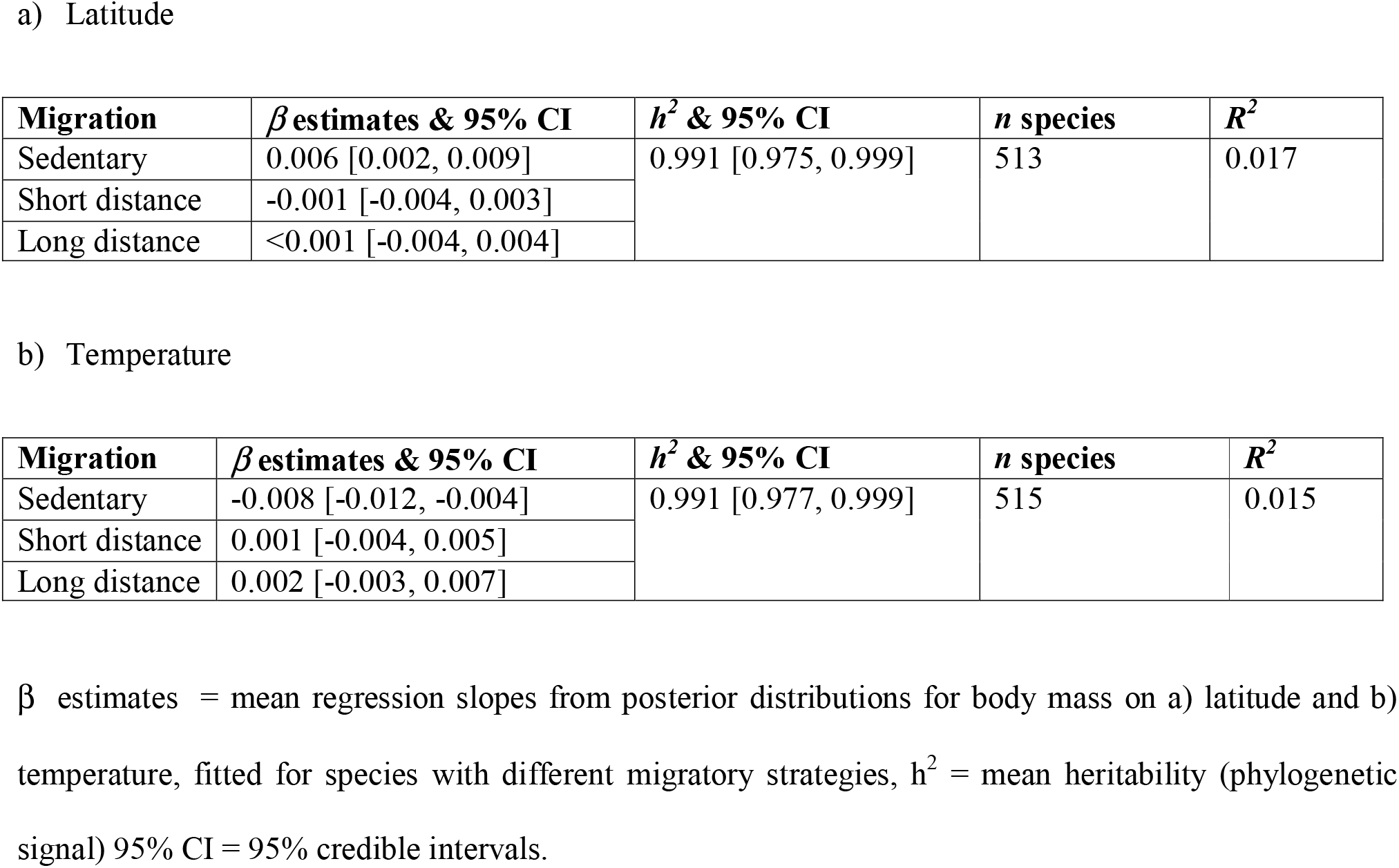
Interaction of Bergmann’s rule with migration.

a) Latitude
| Migration | $\beta$ estimates & 95% CI | $h^2$ & 95% CI | $n$ species | $R^2$ |
| --- | --- | --- | --- | --- |
| Sedentary | 0.006 [0.002, 0.009] | 0.991 [0.975, 0.999] | 513 | 0.017 |
| Short distance | -0.001 [-0.004, 0.003] |  |  |  |
| Long distance | <0.001 [-0.004, 0.004] |  |  |  |

b) Temperature
| Migration | $\beta$ estimates & 95% CI | $h^2$ & 95% CI | $n$ species | $R^2$ |
| --- | --- | --- | --- | --- |
| Sedentary | -0.008 [-0.012, -0.004] | 0.991 [0.977, 0.999] | 515 | 0.015 |
| Short distance | 0.001 [-0.004, 0.005] |  |  |  |
| Long distance | 0.002 [-0.003, 0.007] |  |  |  |

$\beta$ estimates = mean regression slopes from posterior distributions for body mass on a) latitude and b) temperature, fitted for species with different migratory strategies, $h^2$ = mean heritability (phylogenetic signal) 95% CI = 95% credible intervals.

**Figure 1.**
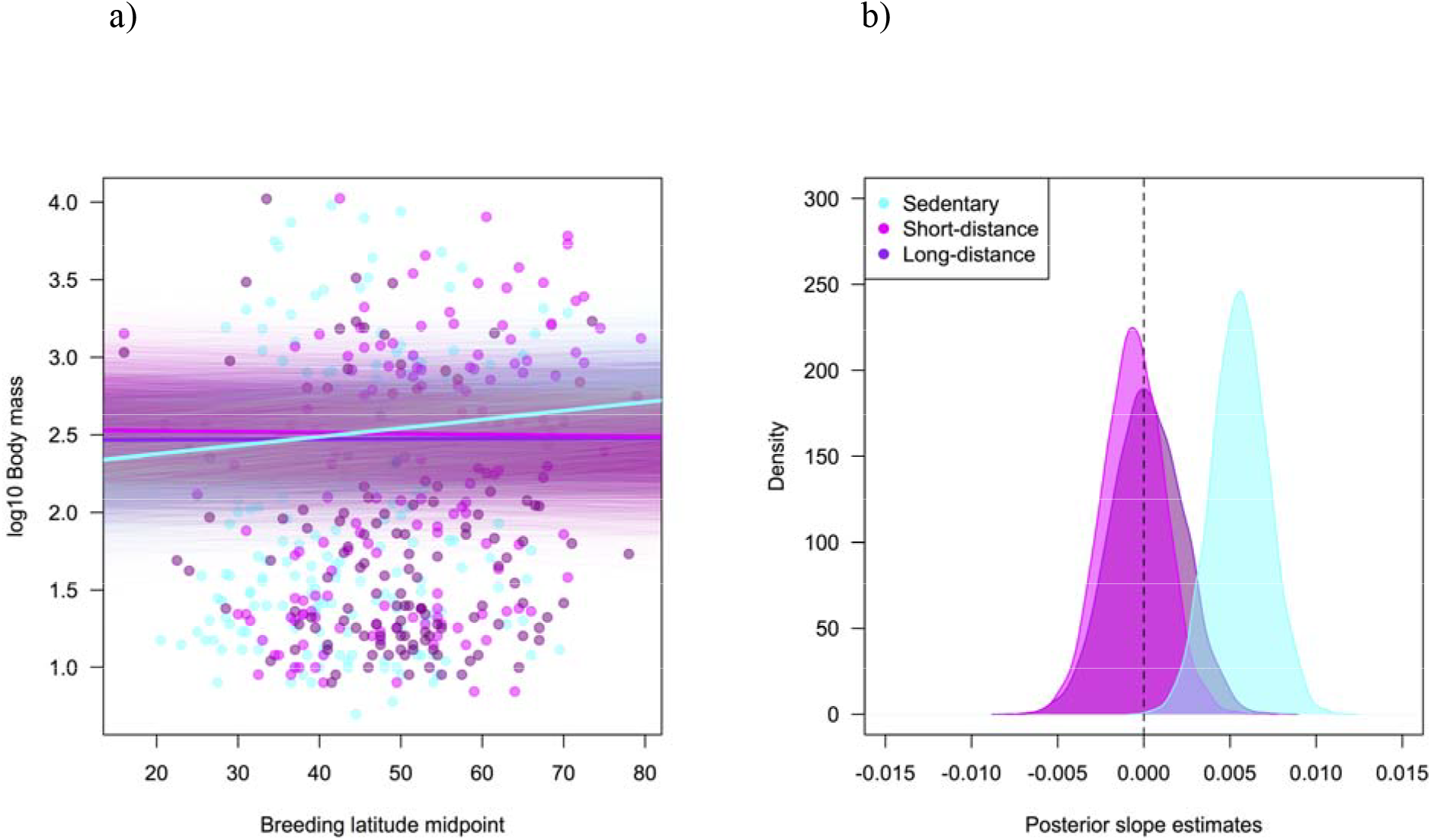
Interaction of Bergmann’s rule with migration. a) Species’ body mass against breeding latitude midpoint with different slopes fitted for sedentary, short-distance migrating and long-distance migrating species. Mean slopes from posterior distributions are indicated by thick lines, while slopes from the entire posterior distributions are plotted as thinner, semi-transparent lines. b) density plot showing posterior distributions of slope estimates for sedentary, short-distance migrating and long-distance migrating species.

When including a three-way interaction between breeding climate, nest type and migration, we find the predicted effects of nest type on conformity to Bergmann’s rule within sedentary species. Among sedentary species, body mass increases with breeding range latitude and decreases with breeding range temperature for open and semi-open nesting species, but not enclosed nesting species. Short- and long-distance migrating species do not conform to Bergmann’s rule at all, regardless of nest type (Table 3, Fig. 2). When re-running analyses separating the effects of nest structure and location, we find similar patterns both for nest structure (Table S5) and location (Table S6).

**Figure 2.**
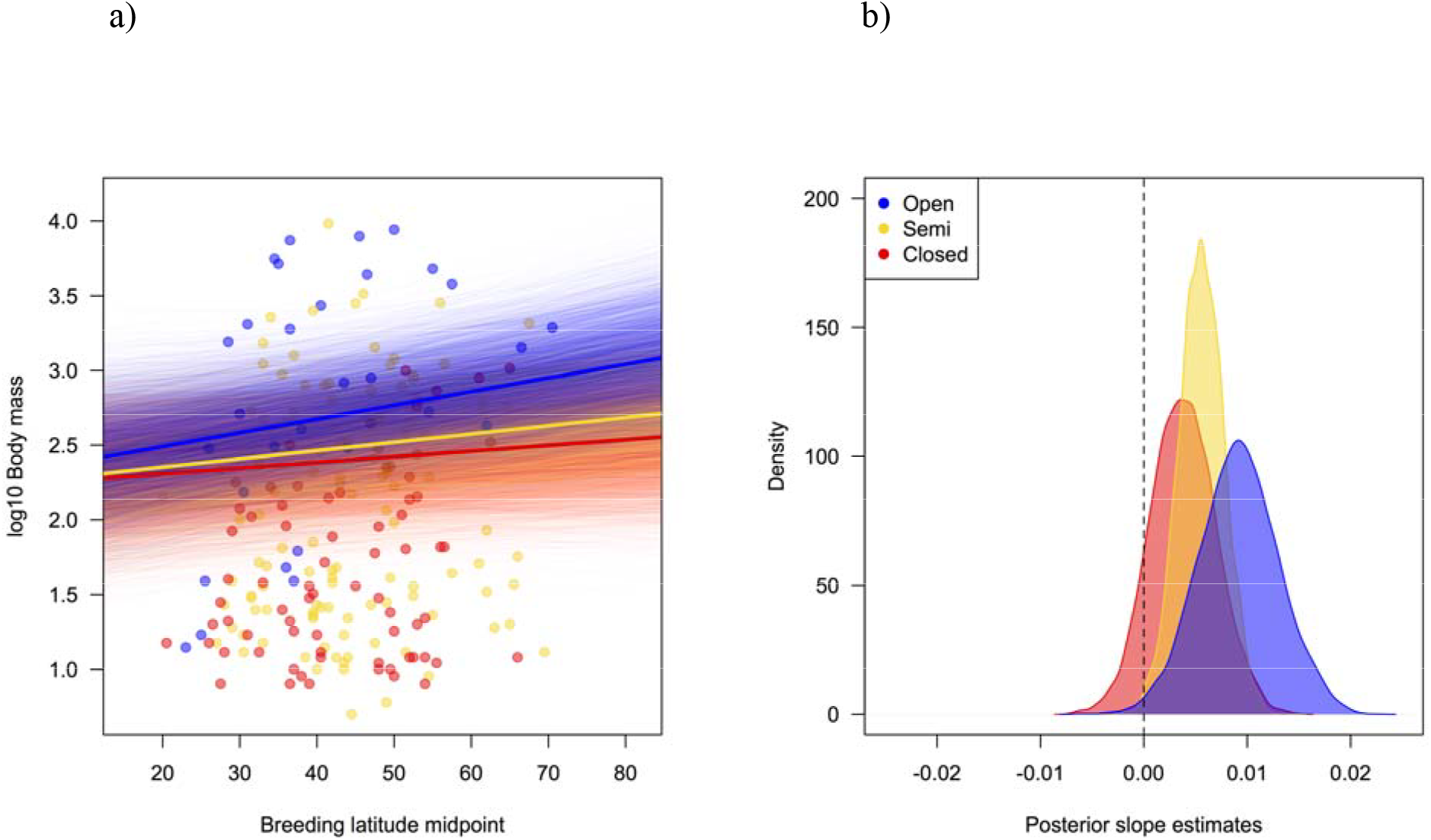
Interaction of Bergmann’s rule with nest design within sedentary species. Species’ body mass against breeding latitude midpoint with different slopes fitted for open, semi-open and enclosed nesting species, within sedentary species only. Mean slopes from posterior distributions are indicated by thick lines, while slopes from the entire posterior distributions are plotted as thinner, semi-transparent lines. b) density plot showing posterior distributions of slope estimates open, semi-open and enclosed nesting species, within sedentary species only.

**Table 3.** Interaction of Bergmann’s rule with migration and nest design.

a) Latitude
| Migration | Nest design | $\beta$ estimates & 95% CI | $h^2$ & 95% CI | $n$ species | $R^2$ |
| --- | --- | --- | --- | --- | --- |
| Sedentary | Open | 0.009 [0.002, 0.016] | 0.988 [0.969, 0.998] | 513 | 0.202 |
|  | Semi | 0.006 [0.001, 0.010] |  |  |  |
|  | Enclosed | 0.004 [-0.003, 0.010] |  |  |  |
| Short | Open | -0.002 [-0.006, 0.004] |  |  |  |
|  | Semi | 0.001 [-0.005, 0.006] |  |  |  |
|  | Enclosed | -0.002 [-0.010, 0.006] |  |  |  |
| Long | Open | <-0.001 [-0.006, 0.006] |  |  |  |
|  | Semi | -0.002 [-0.008, 0.005] |  |  |  |
|  | Enclosed | 0.004 [-0.006, 0.014] |  |  |  |

b) Temperature
| Migration | Nest design | $\beta$ estimates & 95% CI | $h^2$ & 95% CI | $n$ species | $R^2$ |
| --- | --- | --- | --- | --- | --- |
| Sedentary | Open | -0.008 [-0.016, <-0.001] | 0.989 [0.972, 0.999] | 515 | 0.188 |
|  | Semi | -0.009 [-0.014, -0.004] |  |  |  |
|  | Enclosed | -0.006 [-0.014, 0.004] |  |  |  |
| Short | Open | 0.003 [-0.003, 0.010] |  |  |  |
|  | Semi | -0.002 [-0.009, 0.005] |  |  |  |
|  | Enclosed | 0.001 [-0.012, 0.014] |  |  |  |
| Long | Open | 0.003 [-0.004, 0.010] |  |  |  |
|  | Semi | 0.002 [-0.006, 0.011] |  |  |  |
|  | Enclosed | -0.004 [-0.017, 0.010] |  |  |  |

$\beta$ estimates = mean regression slopes from posterior distributions for body mass on a) latitude and b) temperature, fitted for species with different nest designs and migratory strategies, $h^2$ = mean heritability (phylogenetic signal) 95% CI = 95% credible intervals.

## Discussion

We find that conformity to Bergmann’s rule, which predicts that species inhabiting colder environments should have larger body sizes (Bergmann 1847) depends strongly on migration and nest design in Western Palearctic birds. In both long- and short-distance migratory species, body mass is unaffected by climatic conditions in the breeding range, while among non-migrating sedentary species, conformity to Bergmann’s rule is greatest among species whose nests are most exposed to environmental conditions. Our results suggest that smaller bird species can adapt to colder climates either by migrating to avoid extreme winter temperatures, or by breeding in enclosed nests, while colder breeding climates favour larger body sizes in resident, open-nesting species. Therefore, among non-migratory species, enclosed nests may compensate for higher thermoregulatory costs of incubation in smaller-bodied adults breeding in colder climates, thus enabling smaller species to breed in colder climates than their body mass would otherwise allow. Our study is consistent with the idea that birds’ behaviour, and particularly their nesting and migratory strategies, mediate the effect of environmental conditions on species’ morphology.

Among resident species, we find evidence in support of Bergmann’s rule only for species with open or semi-open nests, in which brooding parents are most exposed to adverse weather at northerly latitudes. Therefore, our results support the idea that enclosed nests serve to protect smaller birds against colder climatic conditions at higher latitudes. In additional analyses reported in the Supporting Information, we find the same pattern of results among resident species both for those building open nest structures and siting nests in open locations. Therefore, both enclosed nest structures and open structures built in protected locations can effectively buffer smaller-bodied species against colder conditions at higher latitudes. These results demonstrate the importance of considering not only aspects of nest structure but also of location when investigating ecological correlates of nest design. Our results contrast with a prior study testing the validity of Bergmann’s rule at the intraspecific level within 106 bird species, which found that body mass was no more likely to increase in colder climates within open nesting than enclosed nesting species (Meiri and Dayan 2003). Our findings may differ from those of this study because we test predictions in a larger sample (>500 species) at the interspecific level, capturing far more variation in body mass and climatic conditions. Further, in contrast to prior studies we investigate relationships between body mass and environmental conditions in birds’ breeding ranges specifically rather than across their entire ranges. Nest design is much more relevant to the former since birds’ nests are generally temporary structures built for the purposes of breeding only.

In contrast with our findings, two recent comparative analyses suggest that enclosed nests are protective against exposure to hotter and drier, rather than colder and wetter, climatic conditions. Among diverse geographic regions, the proportion of passerine species with enclosed nests is 2 to 3 times greater in tropical or southern hemisphere regions than in north temperate regions (Martin et al. 2017). Within Australia, meanwhile, the proportion of passerine species building domed nests increases in areas with hotter, drier climates and less vegetative cover (Duursma et al. 2018). Direct comparisons between our results and these prior studies is challenging due to key methodological differences: these studies are based on geographical patterns rather than phylogenetically-informed relationships, and do not incorporate potential interactions with body mass. Discrepancies with our findings may also be partly explained, however, by different approaches to classifying nest types: in contrast to our study, these analyses did not count nests built in cavities as enclosed due to a focus solely on nest structure. In our sample, a substantial proportion (∼20%) of species nest in cavities which may provide effective protection against colder breeding environments in the Northern hemisphere. Taken at face value, however, these differing results suggest that protective effects of enclosed nests against extreme climatic conditions may be region-specific. Enclosed nesting may only have the opportunity to evolve in response to colder climates within the Northern hemisphere, which encompasses far more potential breeding range in temperate and polar climatic zones than the Southern hemisphere.

Along with nest type, we also identify migration as an important mediator of conformity to Bergmann’s rule. Consistent with some previous work at the intraspecific level (Meiri and Dayan 2003, but see Ashton 2002), we find that body mass increases in colder temperatures among sedentary species, but in neither short-distance or long-distance migrants. These findings support the idea that long-distance migrants are less exposed to selection pressures favouring large body size in colder climates as they avoid exposure to the coldest winter temperatures at high latitudes by spending the non-breeding season in warmer environments (Ashton 2002, Meiri and Dayan 2003). Teasing apart the effects of migration and nest type on conformity to Bergmann’s rule, we find that migration has a stronger effect than nest type. While we find predicted effects of nest type within sedentary species, migratory species do not conform to Bergmann’s rule at all, regardless of their nesting behaviour. Therefore, the thermoregulatory benefits of migration override those of nest design, such that enclosed nests provide no additional thermoregulatory benefits for migratory, small-bodied species. This is perhaps unsurprising because migration results in species avoiding extreme winter conditions in the Northern hemisphere altogether, while nest design can only affect exposure to environmental conditions for relatively short periods during breeding. Taken together, our results reveal the interplay between nesting and migration in buffering small-bodied species against cold climates in the Northern hemisphere.

Alternative explanations for our findings may be related to potential systematic changes in the availability of nest sites (Hansell 2000) or food (Martin 1995) over latitudinal gradients: for example, natural cavities and food may be limited in forests at higher latitudes which may mean that smaller and competitively inferior tree cavity-nesting species are prevented from breeding at higher latitudes through competitive exclusion rather than environmental conditions alone. However, these alternatives seem unlikely as natural cavities are not usually in limited supply in the northern hemisphere (Wiebe 2011). Instead, our findings suggest that migration and nest morphology in birds and other animals may help species to breed in climates where they would not necessarily otherwise be able to. Meanwhile the need for streamlined body designs for efficient flight in migrant birds may play a greater role in determining their morphology than conditions on the breeding grounds alone. Nevertheless, these findings are consistent with a prediction of niche construction theory that susceptibility to abiotic selection pressures, such as environmental temperature, can be buffered by species’ alteration of their environments through behaviour, particularly in terms of the location of nesting sites (Odling-Smee et al. 2003). However, since comparative analyses can only identify correlational rather than causal relationships (Nunn 2011), we cannot rule out the possibility of alternative causal explanations. Our results are therefore equally consistent with causal scenarios in which environmental selection pressures drive changes in behaviour rather than vice-versa, or where environmental selection pressures and behaviour influence one another in evolutionary feedback loops. In any case, our findings are significant in that they suggest that fundamental relationships between species’ environments and morphology may be mediated by behaviour.

We have demonstrated that in Western Palearctic birds, body mass increases in colder climates as hypothesised by Bergmann’s rule only in non-migratory species breeding in exposed nests. Our findings are consistent with the idea that migration and enclosed nests compensate for greater thermoregulatory costs in smaller-bodied birds, allowing then to breed in colder environments than expected for their body size. Further research could usefully examine how species’ modification of environments affects responses to environmental selection pressures across more diverse taxa and geographic regions, including across human populations. Our work should also guide future experimental studies on the potential mediating role of nesting and migratory behaviour on the influence of climatic conditions on parental and offspring fitness. We conclude that behaviour, particularly migration, nest-building and nest-site choice, is an important mediator of species’ responses to climatic selection pressures.

## Acknowledgements

We thank Sue Healy and Kevin Laland for useful comments.

## Author contributions

MCM and SES designed the study, MCM collected the data, SES analysed the data and MCM and SES wrote the manuscript.

## SUPPORTING INFORMATION

**Table S1.** Nest design categorisation scheme.

|  | **Open** | **Semi-open** | **Enclosed** |
| --- | --- | --- | --- |
| **Cup** | Open | Semi-open | Enclosed |
| **Plate** | Open | Semi-open | Enclosed |
| **Scrape** | Open | Semi-open | Enclosed |
| **Bed** | Open | Semi-open | Enclosed |
| **Dome** | Enclosed | Enclosed | Enclosed |
| **Dome & tube** | Enclosed | Enclosed | Enclosed |
| **Burrow** | Enclosed | Enclosed | Enclosed |

Scheme used to combine nest structure and location into a single nest design variable. Nest structure (rows) and nest location (columns) categories are combined to form a single nest design variable, capturing differing levels of exposure to environmental conditions influenced by both nest structure and location.

**Table S2.**
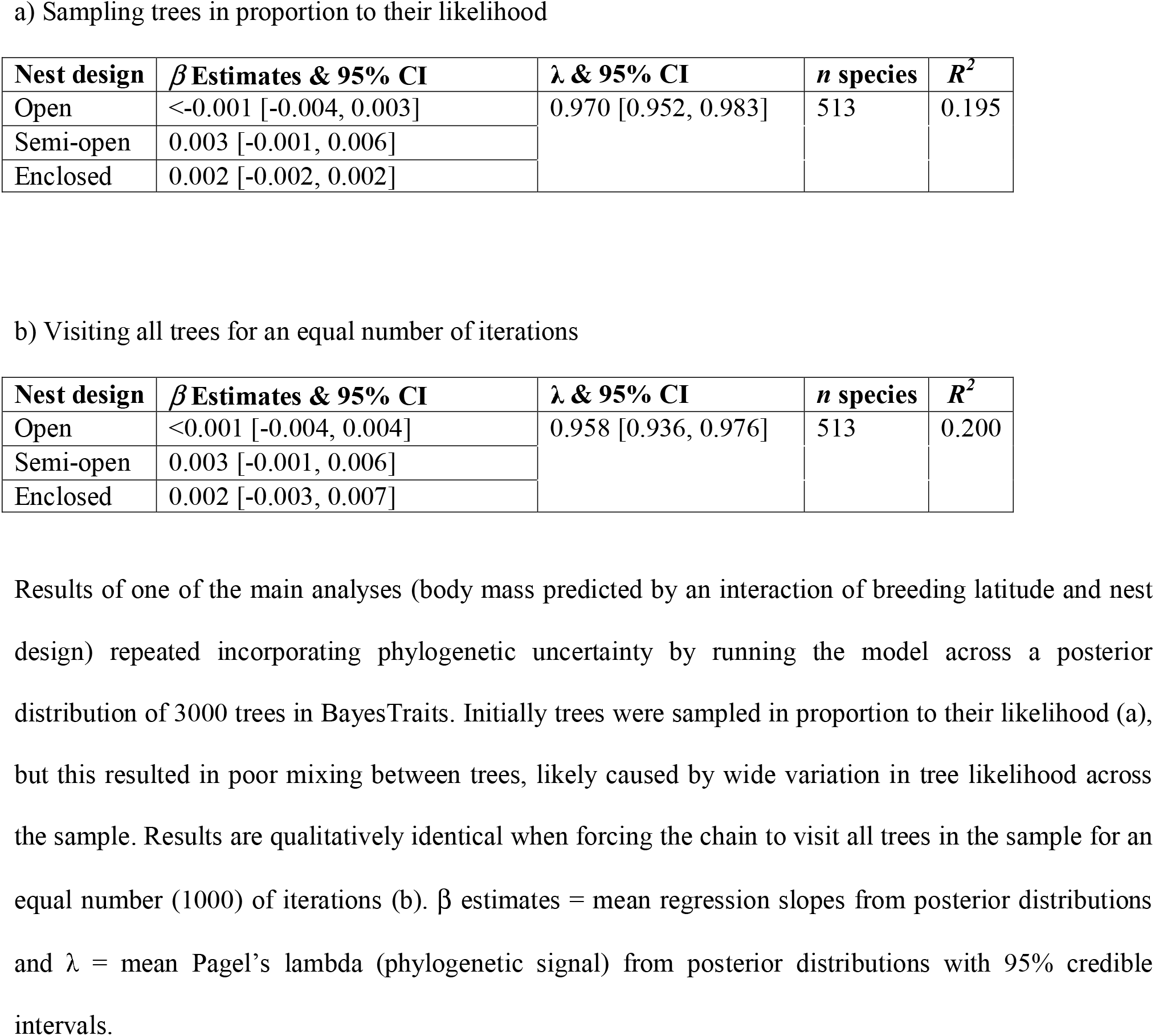
Incorporating phylogenetic uncertainty.

a) Sampling trees in proportion to their likelihood
| Nest design | $\beta$ Estimates & 95% CI | $\lambda$ & 95% CI | *n* species | $R^2$ |
| --- | --- | --- | --- | --- |
| Open | <-0.001 [-0.004, 0.003] | 0.970 [0.952, 0.983] | 513 | 0.195 |
| Semi-open | 0.003 [-0.001, 0.006] |  |  |  |
| Enclosed | 0.002 [-0.002, 0.002] |  |  |  |

b) Visiting all trees for an equal number of iterations
| Nest design | $\beta$ Estimates & 95% CI | $\lambda$ & 95% CI | *n* species | $R^2$ |
| --- | --- | --- | --- | --- |
| Open | <0.001 [-0.004, 0.004] | 0.958 [0.936, 0.976] | 513 | 0.200 |
| Semi-open | 0.003 [-0.001, 0.006] |  |  |  |
| Enclosed | 0.002 [-0.003, 0.007] |  |  |  |

**Table S3.** Bergmann’s rule and nest structure.

| Nest structure | $\beta$ estimates & 95% CI | $h^2$ & 95% CI | $n$ species | $R^2$ |
| --- | --- | --- | --- | --- |
| Open | 0.001 [-0.001, 0.004] | 0.988 [0.971, 0.998] | 513 | 0.055 |
| Enclosed | 0.005 [-0.002, 0.012] |  |  |  |

d) Temperature
| Nest structure | $\beta$ estimates & 95% CI | $h^2$ & 95% CI | $n$ species | $R^2$ |
| --- | --- | --- | --- | --- |
| Open | -0.002 [-0.005, 0.001] | 0.988 [0.970, 0.998] | 515 | 0.050 |
| Enclosed | -0.009 [-0.019, <0.001] |  |  |  |

Results of models allowing the slope of body mass on a) latitude or b) temperature to vary between species with different nest structures (open = cup, plate, scrape or bed, closed = dome, dome and tube or burrow), regardless of location. $\beta$ estimates = mean regression slopes from posterior distributions and $h^2$ = mean heritability (phylogenetic signal) from posterior distributions with 95% credible intervals.

**Table S4.** Bergmann’s rule and nest location.

a) Latitude
| Nest location | $\beta$ estimates & 95% CI | $h^2$ & 95% CI | $n$ species | $R^2$ |
| --- | --- | --- | --- | --- |
| Open | <0001 [-0.003, 0.004] | 0.987, [0.969, 0.998] | 513 | 0.171 |
| Semi-open | 0.003 [<-0.001, 0.006] |  |  |  |
| Enclosed | 0.002 [-0.003, 0.008] |  |  |  |

b) Temperature
| Nest location | $\beta$ estimates & 95% CI | $h^2$ & 95% CI | $n$ species | $R^2$ |
| --- | --- | --- | --- | --- |
| Open | 0.001 [-0.003, 0.006] | 0.989 [0.972, 0.988] | 515 | 0.158 |
| Semi-open | -0.005 [-0.009, -0.002] |  |  |  |
| Enclosed | -0.003 [-0.011, 0.003] |  |  |  |

Results of models allowing the slope of body mass on a) latitude or b) temperature to vary between species with different nest locations (open, semi-open or enclosed), regardless of structure. $\beta$ estimates = mean regression slopes from posterior distributions and $h^2$ = mean heritability (phylogenetic signal) from posterior distributions with 95% credible intervals.

**Table S5.** Interaction of Bergmann’s rule with migration and nest structure.

| Migration | Nest design | $\beta$ estimates & 95% CI | $h^2$ & 95% CI | $n$ species | $R^2$ |
| --- | --- | --- | --- | --- | --- |
| Sedentary | Open | 0.005 [0.002, 0.008] | 0.990 [0.972, 0.999] | 513 | 0.048 |
|  | Enclosed | 0.006 [-0.005, 0.017] |  |  |  |
| Short | Open | <-0.001 [-0.004, 0.003] |  |  |  |
|  | Enclosed | -0.004 [-0.018, 0.012] |  |  |  |
| Long | Open | -0.001 [-0.005, 0.004] |  |  |  |
|  | Enclosed | 0.006 [-0.005, 0.018] |  |  |  |

d) Temperature
| Migration | Nest design | $\beta$ estimates & 95% CI | $h^2$ & 95% CI | $n$ species | $R^2$ |
| --- | --- | --- | --- | --- | --- |
| Sedentary | Open | -0.007 [-0.012, -0.003] | 0.990, [0.973, 0.998] | 515 | 0.046 |
|  | Enclosed | -0.012 [-0.025, 0.002] |  |  |  |
| Short | Open | 0.001 [-0.004, 0.006] |  |  |  |
|  | Enclosed | 0.005 [-0.020, 0.030] |  |  |  |
| Long | Open | 0.003 [-0.002, 0.008] |  |  |  |
|  | Enclosed | -0.009 [-0.024, 0.008] |  |  |  |

Results of models allowing the slope of body mass on a) latitude or b) temperature to vary between species with different migratory strategies (sedentary, short- or long-distance) and nest structures (open = cup, plate, scrape or bed, closed = dome, dome and tube or burrow). $\beta$ estimates = mean regression slopes from posterior distributions and $h^2$ = mean heritability (phylogenetic signal) from posterior distributions with 95% credible intervals.

**Table S6.** Interaction of Bergmann’s rule with migration and nest location.

a) Latitude
| Migration | Nest location | $\beta$ estimates & 95% CI | $h^2$ & 95% CI | $n$ species | $R^2$ |
| --- | --- | --- | --- | --- | --- |
| Sedentary | Open | 0.009 [0.002, 0.016] | 0.988 [0.971, 0.998] | 513 | 0.172 |
|  | Semi-open | 0.006 [0.002, 0.010] |  |  |  |
|  | Enclosed | 0.003 [-0.004, 0.011] |  |  |  |
| Short | Open | -0.002 [-0.006, 0.003] |  |  |  |
|  | Semi-open | 0.001 [-0.004, 0.006] |  |  |  |
|  | Enclosed | <-0.001 [-0.010, 0.010] |  |  |  |
| Long | Open | <-0.001 [-0.006, 0.006] |  |  |  |
|  | Semi-open | -0.002 [-0.008, 0.005] |  |  |  |
|  | Enclosed | 0.003 [-0.009, 0.015] |  |  |  |

b) Temperature
| Migration | Nest location | $\beta$ estimates & 95% CI | $h^2$ & 95% CI | $n$ species | $R^2$ |
| --- | --- | --- | --- | --- | --- |
| Sedentary | Open | -0.008 [-0.016, <0.001] | 0.989 [0.973, 0.999] | 515 | 0.160 |
|  | Semi-open | -0.010 [-0.014, -0.005] |  |  |  |
|  | Enclosed | -0.006 [-0.015, 0.004] |  |  |  |
| Short | Open | 0.004 [-0.003, 0.010] |  |  |  |
|  | Semi-open | -0.002 [-0.009, 0.006] |  |  |  |
|  | Enclosed | -0.001 [-0.015, 0.014] |  |  |  |
| Long | Open | 0.003 [-0.004, 0.010] |  |  |  |
|  | Semi-open | 0.002 [-0.006, 0.010] |  |  |  |
|  | Enclosed | -0.003 [-0.018, 0.012] |  |  |  |

Results of models allowing the slope of body mass on a) latitude or b) temperature to vary between species with different migratory strategies (sedentary, short- or long-distance) and nest locations (open, semi-open or enclosed). $\beta$ estimates = mean regression slopes from posterior distributions and $h^2$ = mean heritability (phylogenetic signal) from posterior distributions with 95% credible intervals.

**Figure S1.**
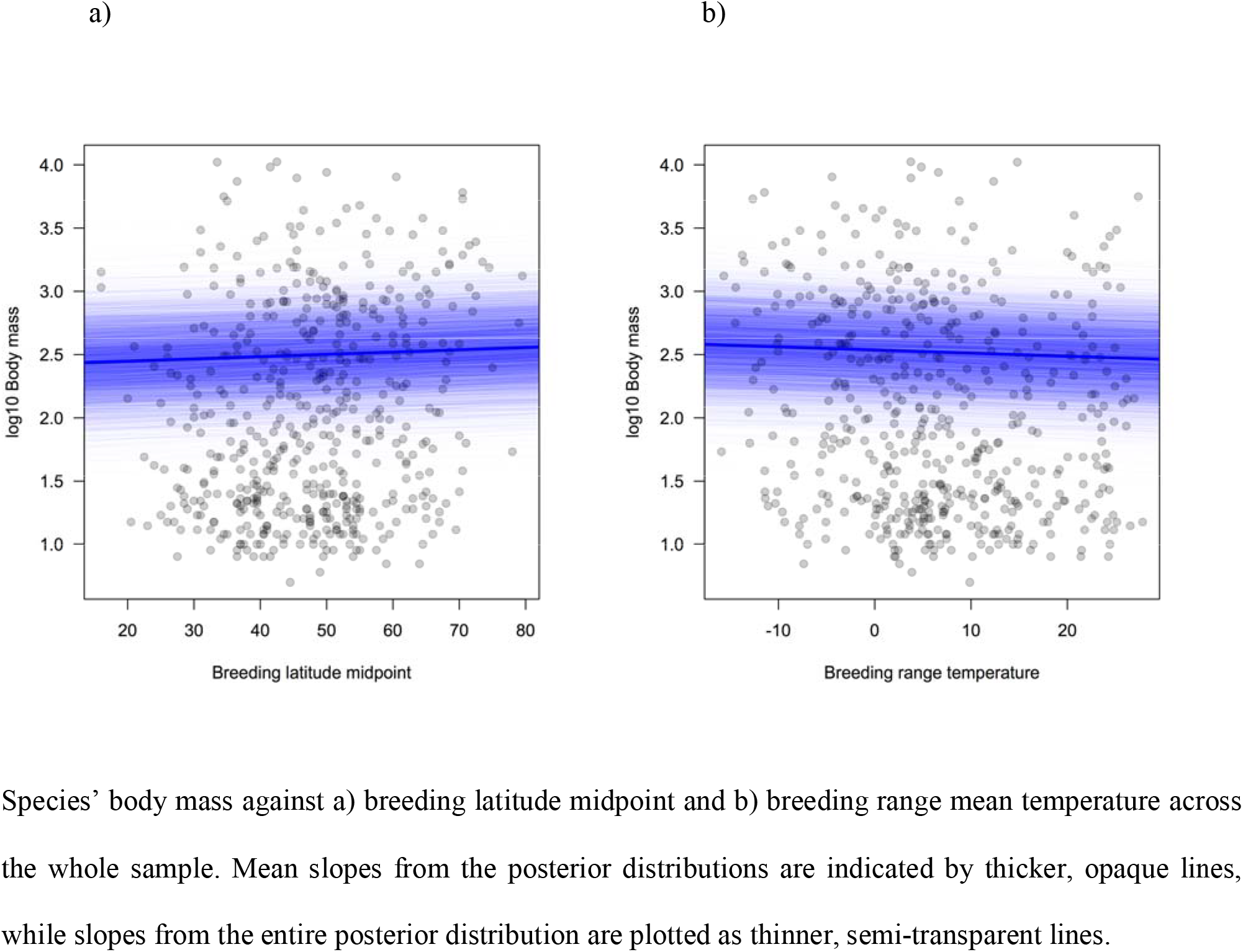
Support for Bergmann’s rule.

